# Anxiety-like behaviors in mice unmasked: Revealing sex differences in anxiety using a novel light-heat conflict test

**DOI:** 10.1101/2022.09.02.506410

**Authors:** Sydney E. Lee, Sung-Hoon Park, John C. Aldrich, Laura K. Fonken, Andrew D. Gaudet

**Affiliations:** Department of Psychology, College of Liberal Arts; Department of Neurology, Dell Medical School; McGovern Medical School, The University of Texas Health Science Center at Houston; Division of Pharmacology and Toxicology, College of Pharmacy

## Abstract

Anxiety and chronic pain afflict hundreds of millions worldwide. Both anxiety and pain are more prevalent in females compared to males. Unfortunately, robust sex differences in human anxiety are not recapitulated in rodent tests, and results from rodent pain studies frequently fail to translate clinically. Therefore, there is a need to develop tests that reflect the differential salience of anxiety or pain-related stimuli between the sexes. Accordingly, here we introduce the Thermal Increments Dark-Light (TIDAL) conflict test. The TIDAL test places an anxiety-relevant stimulus (dark vs. illuminated chamber) in conflict with a heat-related stimulus (incrementally heated vs. isothermic chamber); mice freely explore the dark-heating and illuminated-isothermic chambers. Here, we aim to determine whether the TIDAL conflict test reveals in mice underappreciated sex differences in anxiety and/or heat sensitivity. We establish in four distinct experiments that females on the TIDAL conflict test persist substantially longer on the dark-heated plate, suggesting that females as compared to males exhibit elevated anxiety-like behavior. Mice more strongly prefer the dark plate on the TIDAL conflict test compared to control thermal place preference with both chambers illuminated. We also reveal that an anxiety-relieving drug, paroxetine, reduces mouse preference for the heating dark plate, supporting the validity of the TIDAL test. Therefore, our new TIDAL conflict test reliably unmasks the relative salience of anxiety (vs. heat sensitivity): mice that are female exhibit robust anxiety-like behaviors not consistently observed in classical tests. Future studies should incorporate TIDAL and other conflict tests to better understand rodent behavior and to identify mechanisms underlying anxiety and pain.

## Introduction

Anxiety disorders cost the United States approximately $49 billion annually (Trautmann et al., 2016). Anxiety consists of intense feelings of worry or unease, as well as changes in physical parameters such as blood pressure, and can develop into a debilitating disorder (Dieleman et al., 2016). The 12-month prevalence of anxiety disorders is 11-17%, with a nearly 2x higher prevalence in females (Baxter et al., 2013; McLean et al., 2011; Somers et al., 2006). Females show increased prevalence of separation anxiety, phobias, generalized anxiety, and panic disorders beginning at childhood or adolescence (Altemus et al., 2014; Bekhbat & Neigh, 2018; Donner & Lowry, 2013). Given the high prevalence and burden of these disorders, studying underlying mechanisms and related behaviors could help identify therapies that alleviate maladaptive anxiety.

Anxiety-like behavior is assessed in rodents using validated behavioral assays. For example, the elevated plus maze and the open field test induce a stress response through an aversive event or anticipated aversive event (Bailey & Crawley, 2009), which results in a predictable behavioral output (e.g., thigmotaxis) that is modified based on the rodents prior experience (e.g., predator odor increases thigmotaxis). However, there are several limitations to available test of anxiety-related behavior. First, tests of anxiety-like behaviors in rodents were primarily validated in males to investigate pharmacologic treatments. Current anxiety-related assays detect some sex differences, but results are inconsistent across tests and often diverge from findings in humans (see Börchers et al., 2022; Donner & Lowry, 2013). In rats, females as compared to males travel increased distances and reduce anxiety-like behaviors in the open field test and the elevated plus maze (e.g., Börchers et al., 2022; Johnston & File, 1991; Knight et al., 2021; Scholl et al., 2019). In addition, outcomes regarding rodent sex differences in these anxiety-related tests are inconsistent and fail to recapitulate human sex differences (An et al., 2011; Võikar et al., 2001). A second limitation is that existing tests often evaluate a single variable – e.g., light/dark or enclosed/open space – and therefore might underestimate differences in anxiety-like behavior that would occur in more complex environments. Anxiety-like symptoms in mice of different sexes might appear minimal under baseline conditions, but could be unmasked by placing the anxiety-inducing stimulus in conflict with another factor.

Here, we explore the salience of anxiety vs. heat avoidance in mice using a novel place preference conflict test: the Thermal Increments Dark-Light (TIDAL) conflict test. The conflict test occurs in a place preference apparatus with two chambers connected by a walkway; the conflict test is created by placing a strong “anxiety”-salient stimulus – dark (preferred) vs. illuminated – in conflict with a weaker but increasingly aversive “pain”-relevant thermal choice – increasing heat vs. maintained isothermic temperature. We aim to discover whether the relative salience of anxiety vs. heat sensitivity better reflects anxiety-related sex differences reported in ethological and clinical settings. Both anxiety-like behaviors and heat sensitivity are expected to be increased in female compared to male mice, so it is unclear which stimulus would be more salient to mice in our newly designed conflict test. The TIDAL conflict test is used here to test whether sex affects the salience of anxiety vs. heat avoidance. We find that the TIDAL conflict test does not clearly expose group differences in pain-related heat sensitivity; rather, the TIDAL conflict test is a compelling, reliable, and valid tool for unmasking previously underappreciated differences in anxiety-like behavior.

## Materials and Methods

### Animals and housing

All housing, care, and testing were approved by The University of Texas at Austin Institutional Animal Care and Use Committee. All animals were fed standard chow and filtered tap water *ad libitum* and maintained on a 12:12 light/dark cycle. Adult (8-12 weeks old) male and female C57BL/6J mice (Jackson stock 000664) were tested during the light cycle. Mice were housed in pairs. Mice in all treatment groups were numbered randomly to ensure researchers were blind to group. At the experimental endpoint, mice were injected with an overdose of Pentobarbital (200-270 mg/kg, MWI Animal Health 011355) and tissue was collected for potential later analyses.

### Behavioral tests for anxiety-like behavior

#### Thermal Increments Dark-Light (TIDAL) Conflict Test

The TIDAL conflict apparatus is a modified thermal place preference (TPP) apparatus (Ugo Basile, Cat. No. 35250), which consists of two cylinders (20 cm diameter x 25 cm high) connected by a narrow center walkway (**Fig. 1**). For TIDAL testing, one cylinder (the “light chamber”) is kept in constant light and at a temperature of 31°C, which is an isothermic temperature for mice; in contrast, the other cylinder (“dark chamber”) is covered with a fitted opaque lid and a flexible opaque outside cover to maintain darkness inside the cylinder and the temperature is manipulated from 31 to 44°C (**Fig. 1A,D**). Room illumination levels were 1200 lux, light chamber illumination levels were 1000 lux, and dark chamber illumination levels were 8 lux. The Ugo Basile device is costly; other laboratories considering similar studies could use the Ugo Basile device (which has plates that distribute temperature evenly); explore other commercially available options; or create custom thermal preference chambers using hot plates. In addition, the center walkway was covered with a clear plastic film “roof” to limit mouse interest in escaping through the open space. Mice are not acclimated to the apparatus prior to testing. Pilot studies used a wide range of temperatures (31-52°C) and times at each temperature (5-10 min) (**Fig. 1C**). Based on these pilot studies, to optimally detect the salience of anxiety vs. thermal avoidance we defined the following parameters: Prior to testing, mice are brought into the behavioral testing room and allowed to acclimate in their homecage for ∼30 minutes. Following room acclimation, mice are placed on the illuminated side in TIDAL or the equivalent side in the TPP, and initially allowed to explore the apparatus for 5 minutes with both plates at 31°C (exploratory phase; initial light-dark test); next, an additional five minutes is spent with both plates at 31°C; then, the temperature on the dark plate is raised to 39°C and increased by 1°C every five minutes to a maximum temperature of 44°C (with the light plate maintained at an isothermic 31°C). Thus, a single mouse completes the TIDAL conflict test (or control thermal place preference) assay in 40 minutes (Fig. 1D).

**Figure 1.**
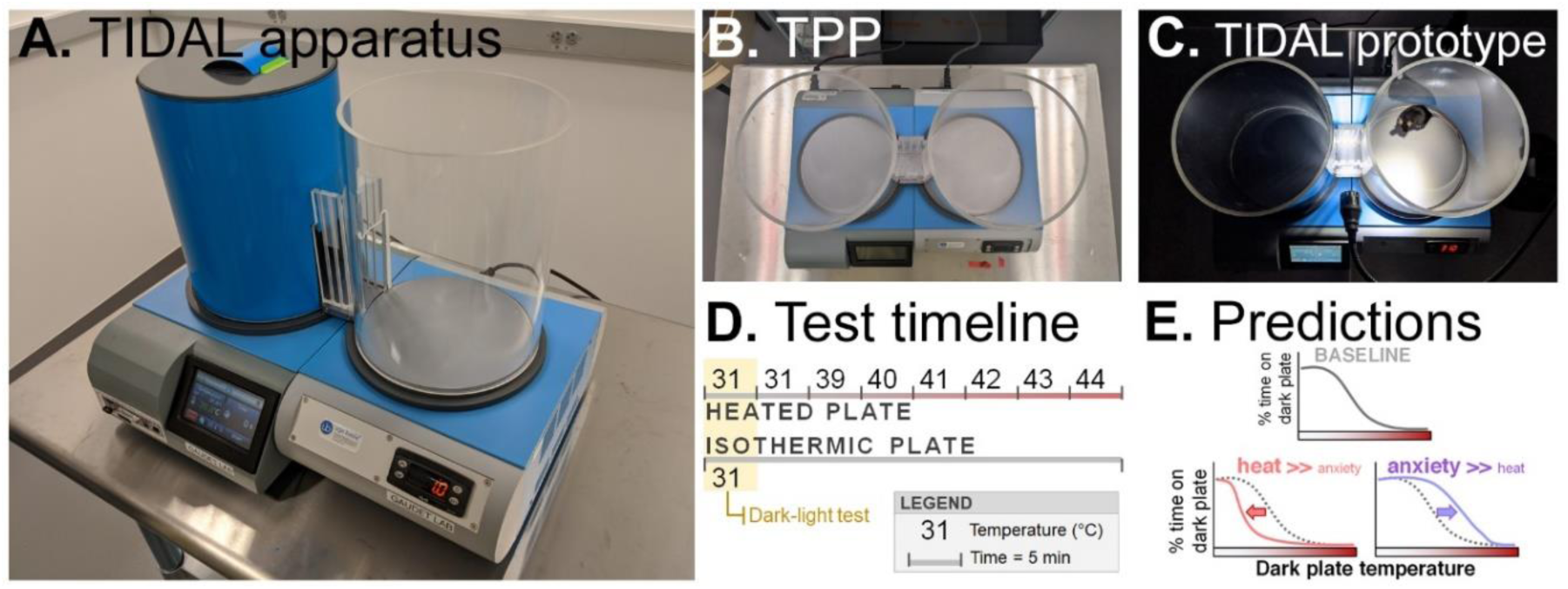
The Thermal Increments Dark-Light (TIDAL) conflict test for exploring anxiety-like behavior and thermal sensitivity in mice: design, controls, timeline, and predictions. **A.** Final, optimized setup of the TIDAL apparatus. Note that the overhead room lights are on, and the dark-heated plate is covered with a lid and opaque film. **B.** Setup of the Thermal Place Preference (TPP) apparatus, a control condition that removes the light-dark (anxiety-related) aspect of the TIDAL conflict test to isolate the effects of shifting heat on one plate. **C.** TIDAL prototype setup for data described in Fig. 2. Note lighting using focal lamps (rather than overhead lights) and lack of a lid on the top of the dark plate. **D.** Optimized TIDAL/TPP test timeline. In TIDAL, the heat-shifting plate is dark and the isothermic plate is lighted, whereas in TPP both plates are lighted. Mice are placed in the apparatus with both sides set to 31°C. The first 5 min are recorded as an initial dark-light test, a control condition that reveals baseline preference for the dark (a measure of anxiety-like behavior). Mice remain in the apparatus as the heating plate continues for 5 min at 31°C, then 39°C, then increases incrementally 1°C every 5 min through 44°C. The lighted isothermic side is maintained throughout at 31°C. **E.** Using the TIDAL conflict test, mice are expected to initially prefer the dark side, but will increasingly avoid the dark plate as its temperature increases. We predict that sex and neurotrauma will alter the salience of heat discomfort vs. anxiety-related sensitivity, manifesting as shifted dark plate preferences over time.

#### Thermal Place Preference (TPP) Assay

The TPP assay is used as a control to isolate thermal sensitivity from the anxiety-like portion of the TIDAL conflict assay. The TPP setup is the same as the TIDAL setup (two cylinders connected by a center walkway), except that both the heated side and the side maintained at 31°C are exposed to room lighting (Fig. 1B) – i.e., the heated side is lighted, not dark as in the TIDAL conflict assay. Next, the same incremental temperature increases are applied.

#### TIDAL and TPP – testing, automated video recording, and analysis

Mice tested on TPP and TIDAL assays were interspersed throughout the day (i.e., during the light phase – Zeitgeber time 1-11). Unless otherwise noted, distinct mice were used for these tests to avoid effects of learning observed in repeated testing. The percent time spent in the dark cylinder (dark cylinder time)/(dark + illuminated time) – center walkway time excluded), distance traveled, and dark crossings were automatically recorded and scored using an overhead video camera and EthoVision software. EthoVision is an applied video tracking software capable of real-time analysis of mouse behavior, movement, and activity at the pixel-level. Time in the center walkway was excluded from analyses in the main manuscript for two reasons: (1) the surroundings in the center zone differed from the test chambers; and (2) analyzing behavior in the identically-shaped illuminated chamber vs. dark-heating chamber enabled a two-chamber preference comparison with equal preference clearly defined at 50% time in each chamber. Including the center zone in analysis had little effect on the percent dark plate preference differences between groups. The arena was cleaned with 70% ethanol between trials.

#### Paroxetine administration

Paroxetine hydrochloride hemihydrate was obtained from Sigma-Aldrich and diluted in vehicle (mixture of 10% DMSO and 5% TWEEN 80 in saline) before experiments. Mice were restrained and administered 10 mg/kg I.P. paroxetine 1 hour prior to TIDAL conflict testing. After paroxetine administration, mice were placed into a novel holding cage for 1 hour to mitigate injection stress.

## Experiments and mouse numbers

Unless noted otherwise, all mice were 8-12 weeks old at time of testing. In Experiment 1 (initial characterization of TIDAL conflict test with broad temperature range), groups included female-TIDAL (n=13) and male-TIDAL (n=15). In Experiment 2 (sex comparison on optimized TIDAL vs. TPP), groups included female-TPP (n=12), female-TIDAL (n=14), male-TPP (n=13), and male-TIDAL (n=15). In Experiment 3 (sex differences in TIDAL over two sessions), groups included female (n=6; 1 excluded from session 2 [EthoVision analysis detection issues]) and male (n=6) mice. In Experiment 4 (TIDAL with paroxetine) groups included female-vehicle (n=8), male-vehicle (n=8), female-paroxetine (n=9; 1 excluded [EthoVision analysis detection issues]), and male paroxetine (n=11).

## Statistics

Mouse TIDAL behavior (dark plate preference, distance traveled) was analyzed using one-, two-, or three-way ANOVA (repeated measures, when appropriate), followed by Bonferroni *post-hoc* tests. In experiments with two groups, a Student’s *t-*test (or nonparametric Mann–Whitney *U* test) was performed. Prism 9 (GraphPad) was used for visualizing data and SigmaPlot 14 (SPSS) was used for statistical analyses.

## Results

### Placing anxiety in conflict with incrementally increasing temperature unmasks sex differences in the salience of anxiety (vs. heat sensitivity)

Women have increased susceptibility to anxiety compared to men, yet mouse models of anxiety show mixed results between the sexes. Women also withdraw more quickly from painful heat stimuli (Bartley & Fillingim, 2013; Bragdon et al., 2002; Feine et al., 1991; Reddan et al., 2020; Rhudy & Meagher, 2001). To explore whether differences in anxiety-vs. pain-like behavior in mice could be uncovered by incorporating conflicting stimuli, we developed and evaluated the TIDAL conflict test.

In our first experiment, individual female or male mice were placed on the TIDAL apparatus (trial setup as in Fig. 1C – initial test with lighting using directed light, rather than existing room overhead lighting, and no top on the dark plate). Temperature on the illuminated plate was maintained at an isothermic 31°C, whereas the temperature on the dark plate was increased incrementally: dark plate temperature started at 31°C (acclimation – dark-light test), then maintained at 31°C for another five minutes, then raised incrementally through 42, 44, 46, 48, 50, and 52°C (five minutes at each temperature) (Fig. 2). If females as compared to males had increased salience of heat hypersensitivity, we would expect them to avoid the heated-dark plate at lower temperatures (curve shifted left); conversely, if females had increased salience of anxiety, we predict they would persist on the heated-dark plate to higher temperatures (curve shifted right) (Fig. 1E).

**Figure 2.**
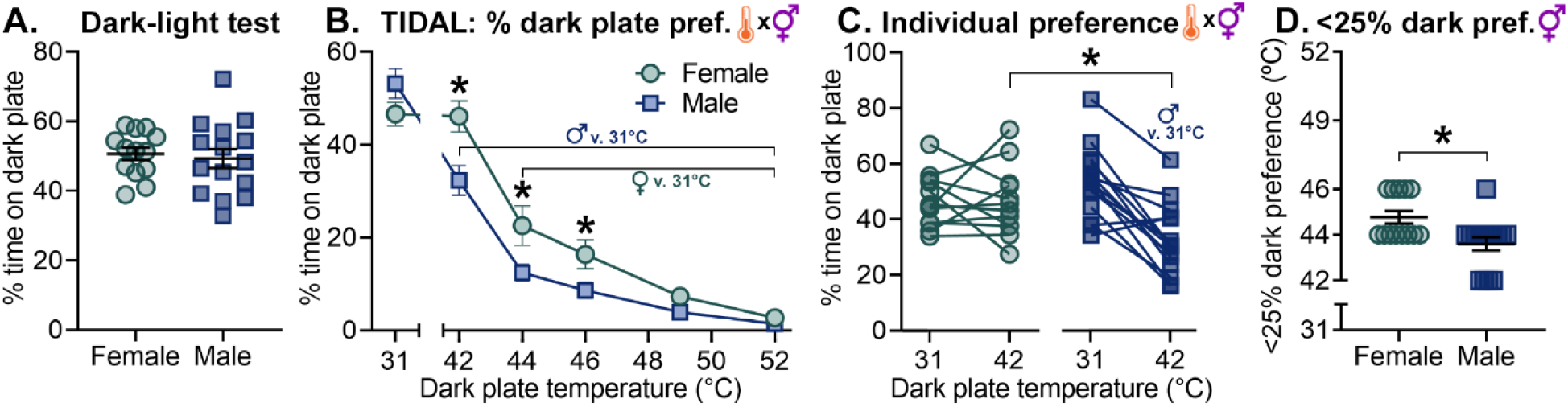
An initial study using the Thermal Increments Dark-Light (TIDAL) conflict test between 42-52°C showed that females as compared to males displayed increased salience of anxiety. **A.** In the dark-light test with both plates at 31°C, females and males showed no significant difference in dark plate preference. **B.** TIDAL test with increasing temperature on the dark plate only. Compared to males, females showed extended preference for the heating dark plate (vs. the constant 31°C lighted plate). **C.** Dark plate preferences of individual mice with the dark plate at 31°C and 42°C. At 42°C, females averaged 46% time on the dark plate, whereas males averaged 32% dark plate preference. **D.** Threshold at which female and male mice showed less than 25% preference for the dark plate. *n*=13 female, *n*=15 male mice; * indicates *p* < 0.05 between female and male mice; “thermometer x gender” symbol indicates significant temperature x sex interaction; gender symbol alone indicates main effect of sex ;“female/male symbol v. 31°C” indicates within-sex difference vs. baseline 31°C.

On the dark-light test, females and males spent a similar percentage of time on the dark plate (Fig. 2A) (females: 51 ± 2%, males: 49 ± 3%; *t*-test, *p* > 0.05). The dark plate preference on the dark-light test was surprisingly low, so in subsequent studies we refined the dark-light contrast between chambers (see below).

In the TIDAL conflict test, temperature increases caused mice to spend progressively less time on the heated dark plate. Compared to baseline preference at 31°C, males spent less time on the dark plate beginning at 42°C, and females had reduced dark plate preference beginning at 44°C (males: 31°C: 53 ± 2%, 42°C: 32 ± 2%; *p* < 0.001) (females: 31°C: 47 ± 3%, 44°C: 23 ± 3%, *p* < 0.001) (two-way RM ANOVA with Bonferroni; sex x temperature interaction *F*_5,130_ = 5.13; *p* < 0.001) (Fig. 2B). Both females and males spent <10% of time on the dark plate at 49 and 52°C, suggesting that these temperatures were strongly aversive.

This TIDAL conflict test exposed notable sex differences in salience of anxiety: females as compared to males spent more time on the dark plate at 42, 44, and 46°C (Fig. 2B). At 42°C females spent 44% more time than males on the heated dark plate (female 42°C: 46 ± 3%, male 42°C: 32 ± 2%; two-way RM ANOVA, *p* < 0.001), and males, but not females, reduced dark plate preference at 42°C (vs. 31°C) (Fig. 2C). Further, females spent <25% of time on the dark plate at a higher temperature than males (Mann-Whitney Rank Sum test, *p* < 0.05) (Fig. 2D). Females and males in the TIDAL test covered a similar distance and females modestly increased percent crossings into the dark plate area (**Fig. S1**). Female and male mice showed surprisingly low baseline dark plate preference (∼50%) and a sharp decrease in time spent on the dark plate with these temperature increases; thus, for subsequent experiments, the TIDAL conflict test was further optimized as outlined below. Overall, this initial study suggests that amplified anxiety-like behavior in female mice can be exposed by placing a known anxiety-related stimulus (dark vs. light) in conflict with increasing temperature.

### Refining the TIDAL conflict test temperature window further illuminates that female mice compared to male mice exhibit increased salience of anxiety

To further optimize the TIDAL conflict test, we improved light-dark contrast in the apparatus (by adding a lid and using overhead lights; Fig. 1A) and refined the temperature range used to better capture the salience of anxiety vs. heat aversion (31°C for 10 min; then 39-44°C, with 1°C increases every 5 min; Fig. 1D). These lighting and temperature range improvements aimed to increase resolution for detecting group differences in TIDAL preferences.

On the dark-light test, females and males undergoing TIDAL both preferred the dark plate vs. the illuminated plate (Fig. 3A) (females: 70 ± 5%, males: 61 ± 3%; *t*-test, *p* > 0.05), suggesting that our apparatus refinements improved light-dark contrast between the two plates. As expected, female and male mice in the control TPP assay – with both plates identically illuminated – had no significant difference in preference for the equivalent-but-illuminated plate (females: 58 ± 4%, males: 58 ± 4%; *t*-test, *p* > 0.05) (data not shown). Here, the dark-light test did not expose significant differences in dark plate preference between females and males.

**Figure 3.**
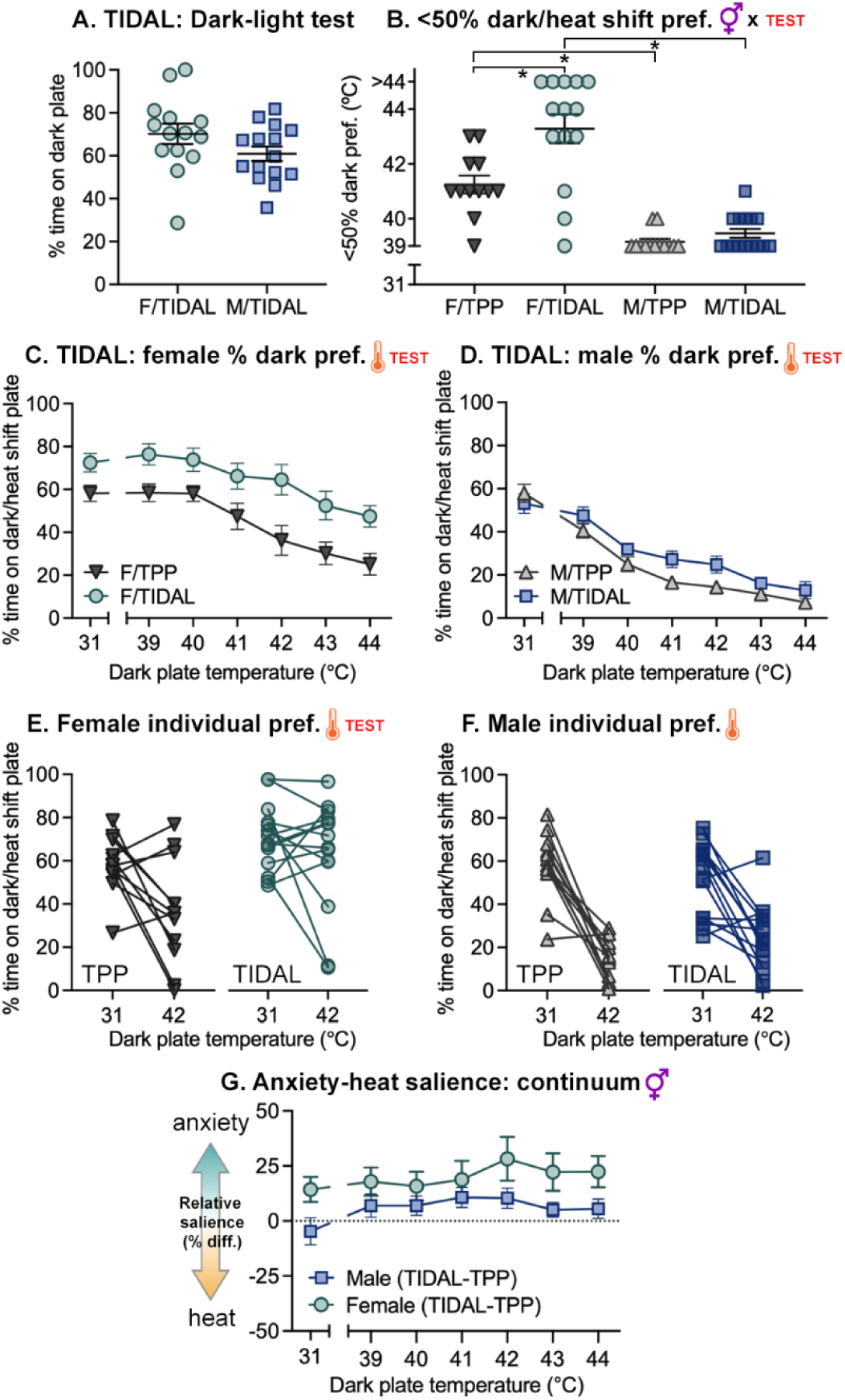
The TIDAL conflict test exposes differences in anxiety-like behavior at optimized incremental heat shift range (39-44°C). Adult female and male mice were tested on the TIDAL conflict test (heated side dark; isothermic side lighted) or on the thermal place preference (TPP) test (both sides lighted). **A.** In the dark-light test with both plates at 31°C, females and males showed similar dark plate preferences. **B.** Threshold at which female and male mice showed less than 50% preference for the dark or heat shift plate in the TIDAL conflict test and TPP test with increasing temperature on the dark/heat shift plate only. Female/TIDAL mice increased <50% threshold temperature compared to both female/TPP and male/TIDAL mice. **C,D.** Female (**C**) and male (**D**) TIDAL/TPP test with increasing temperature on the dark/heat shift plate only. Compared to TPP mice, female and male mice tested on the TIDAL conflict test had heightened preference for the heating dark plate vs. the constant 31°C lighted plate. **E-F.** Dark/heat shift plate preferences of individual mice with the dark/heat shift plate at 31°C and 42°C. At 42°C, TPP females averaged 36% time on the lighted-heated plate, whereas TIDAL females averaged 65% preference for the dark-heated plate (**E**). At 42°C, TPP males averaged 14% time on the lighted-heated plate, whereas TIDAL males averaged 25% preference for the dark-heated plate (**F**). **G.** Anxiety-heat salience continuum. Difference scores were calculated to better delineate differences between TIDAL and control TPP groups. Subtracting control TPP from TIDAL percent dark plate preference, female mice showed increased salience of anxiety-inducing stimulus (dark) vs. heat relative to males. n=12 TPP female, n=14 TIDAL female, n=13 TPP male, n=15 TIDAL male mice; * indicates p < 0.05 between female and male mice; “gender x TEST” symbol indicates significant sex x test interaction; thermometer or TEST symbols alone indicate significant main effects of temperature and TPP/TIDAL, respectively.

Next, behavior on the TPP (control) vs. TIDAL conflict test was assessed in female vs. male mice. Female TIDAL mice remained in the dark portion of the apparatus at higher temperatures than males, suggesting that females exhibit amplified anxiety-like behavior (Fig. 3B**-F**). Indeed, females as compared to males preferred the heated plate to higher temperatures in both tests (Fig. 3B), and mice of both sexes remained on the heated plate longer under TIDAL conditions (with the heated plate also being dark) (Fig. 3C**,D**). Female mice in TIDAL, compared to female-TPP mice, more strongly preferred the heated plate throughout the temperature increases (two-way RM ANOVA; main effect of test [*F*_1,144_ = 11.31] and temperature [*F*_6,144_ = 21.27]; both *p* < 0.005) (Fig. 3C). Similarly, male mice tested on TIDAL (vs. TPP) persisted on the heated plate to higher temperatures (two-way RM ANOVA; main effect of test [*F*_1,156_ = 4.097, *p* = 0.05] and temperature [*F*_6,156_ = 61.71, *p* < 0.001]) (Fig. 3D). Male TPP mice reduced preference for the heat shift plate earlier compared to female TPP mice and compared to male TIDAL mice. Focusing on a pivotal temperature, we found at 42°C that females in particular preferred the heated-dark TIDAL plate vs. the illuminated TPP heated plate (female-TPP 42°C: 36 ± 6%; female-TIDAL 42°C: 65 ± 6%; two-way RM ANOVA, main effects of both temperature and test, *p* < 0.05), whereas male dark plate preference at 42°C was not significantly different between TIDAL and TPP mice (male-TPP 42°C: 14 ± 4%; male-TIDAL 42°C: 25 ± 4%; *p* > 0.05; main effect of temperature only) (Fig. 3E**,F**). Indeed, Therefore, using TPP vs. TIDAL to establish whether eliminating light from the heating chamber extends the duration of preference for the heated plate, our results suggest that the TIDAL conflict test can be used in female and male mice to assess anxiety-like symptoms.

Comparing between sexes, female TPP and TIDAL mice showed higher heat-shift plate preference than males on those same tests (Fig. 3A**-D**). Further, using difference scores to summarize the data, females on TIDAL compared to TPP increased preference for the heated-dark side compared to males (Fig. 3G). Females traveled further per minute in the illuminated area and crossed into the dark chamber more frequently (**Fig. S2**). Overall, these data suggest that this temperature range is well-suited to assess salience of anxiety vs. heat, and that the test effectively unveils anxiety-like behavior (TIDAL vs. TPP results). In addition, females as compared to males strongly increased anxiety-like behavior in TIDAL to an extent that is not consistently observed in a simple dark-light test.

### Over two repeated sessions, females as compared to males maintain prolonged dark plate preference under hyperthermic conditions

Next, we aimed to replicate our sex differences in TIDAL conflict test behavior in a separate cohort to establish reproducibility of our results, and we sought to determine whether mice with prior exposure to TIDAL show evidence of learning.

To address this, a cohort of female and male mice completed two identical TIDAL conflict test sessions separated by two weeks. In the dark-light test over two sessions, females as compared to males had higher preference for the dark plate (two-way RM ANOVA, main effect of sex *F*_1,9_=14.58, *p* < 0.005) (Fig. 4A). This sex difference was particularly notable in Session 2, when females exhibited 83% dark plate preference vs. males’ 54% dark preference. Together, these dark-light test data suggest that females on the dark-light test exhibit anxiety-like behavior, and that the sex difference in anxiety-like behavior is exaggerated in a second exposure to the dark-light apparatus.

**Figure 4.**
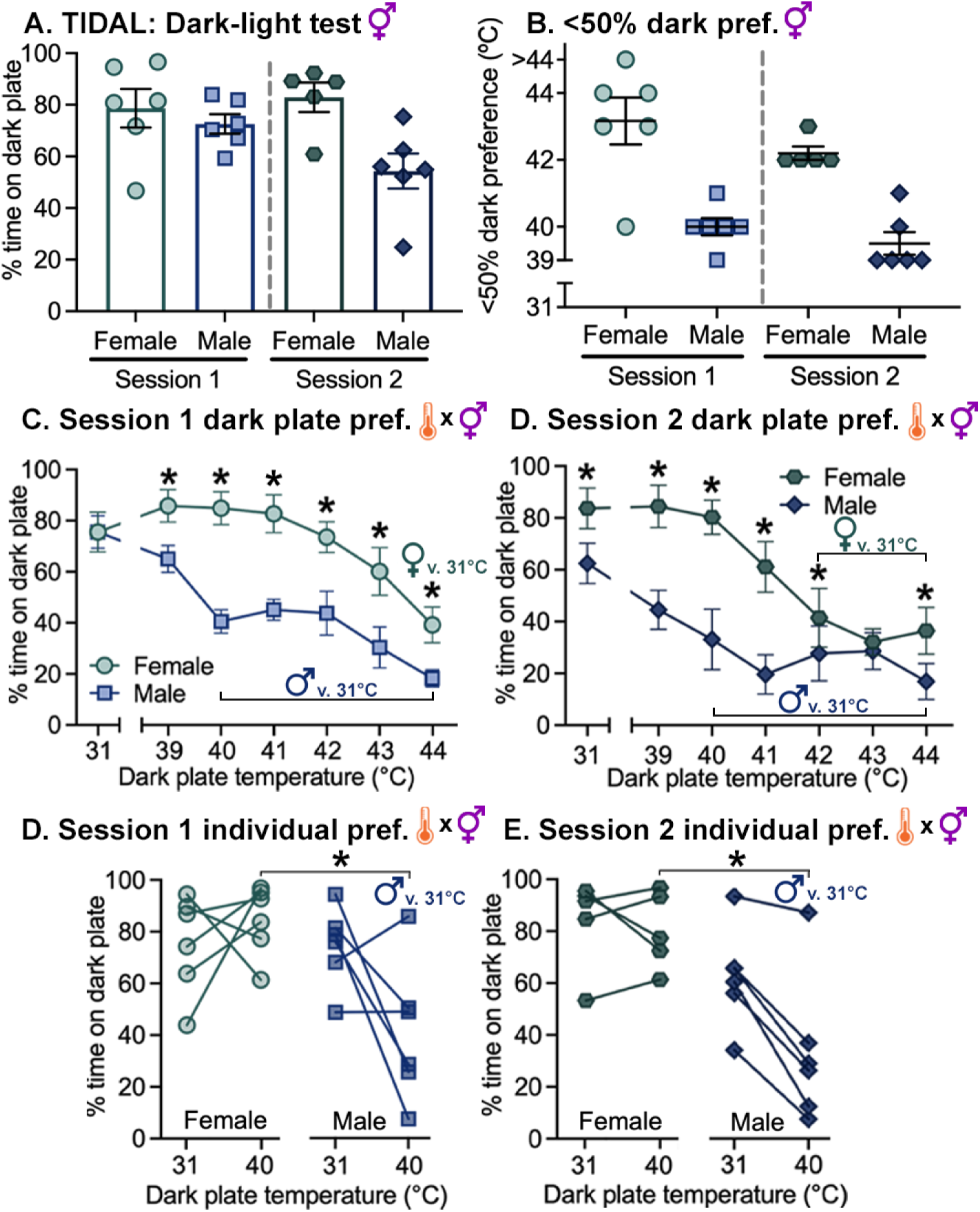
Sex differences in TIDAL conflict test behavior persist across two repeat sessions. Female and male mice completed the TIDAL conflict test twice, two weeks apart. A. In Session 1 and 2 dark-light tests with both plates at 31°C, females had increased dark plate preference. B. Threshold at which mice showed less than 50% preference for the dark plate; females persisted on the heated-dark plate longer than males. C. In the Session 1 TIDAL conflict test, mice of both sexes reduced time spent on the dark plate as temperatures increased. Further, female as compared to males mice preferred the heated-dark plate at all hyperthermic temperatures tested (39-44°C). D. In the Session 2 TIDAL conflict test, mice of both sexes reduced time spent on the dark plate as temperatures increased more quickly than in Session 1. Female as compared to males mice preferred the heated-dark plate at all hyperthermic temperatures tested except 43°C. E. Session 1 dark plate preferences of individual mice with the dark plate at 31°C and 40°C. At 40°C, females had higher dark plate preference than males (F: 85%; M: 42%). F. Session 2 dark plate preferences of individual mice with the dark plate at 31°C and 40°C. At 40°C, females had higher dark plate preference than males (F: 80%; M: 33%). Session 1: *n*=6 female mice, *n*=6 male mice; Session 2: *n*=5 female mice, *n*=6 male mice. * indicates *p* < 0.05 between female and male mice; “thermometer x gender” symbol indicates significant temperature x sex interaction; gender symbol alone indicates significant main effect of sex.

Next, we assessed TIDAL conflict test behavior in both sexes over two sessions. First, we compare TIDAL results by sex: TIDAL females as compared to males had increased preference for the heated-dark plate in both Sessions 1 and 2 (Fig. 4), thereby closely recapitulating results from our previous studies (in Fig. 2**, 3**). In both TIDAL sessions, females spent <50% of their time on the dark plate at higher temperatures compared to males. (Sessions 1 and 2: two-way RM ANOVA, main effect of sex, *p* < 0.05) (Fig. 4B). In TIDAL Sessions 1 and 2, female as compared to males mice increased preference for the heat shift-dark plate from 39-44°C (Session 1: two-way RM ANOVA with Bonferroni, sex x temperature interaction, *F*_6,60_=6.22, *p* < 0.001, female vs. male between 39-44°C) (Session 2: sex x temperature interaction, *F*_6,54_=4.04, *p* < 0.005, female vs. male between 39-42°C and 44°C) (Fig. 4C**,D**). One key temperature that exposed sex differences was 40°C: in Session 1 at 40°C, dark plate preference was 85% for females and 41% for males, and Session 2 dark plate preference was 80% for females and 33% for males, respectively (Fig. 4E**,F**). Thus males, but not females, reduced their preference for the dark plate at 40°C (vs. same sex at 31°C), and males had lower preference at 40°C vs. females (Sessions 1 and 2: two-way RM ANOVA with Bonferroni, sex x temperature interactions, both *p* < 0.05). In addition, females more frequently crossed into the dark chamber (**Fig. S3**). These results bolster our previous findings, showing that females as compared to males exhibit robust anxiety-like behavior in the TIDAL conflict assay with enhanced willingness to remain on an aversive temperature stimulus to avoid an illuminated chamber. Further, these data reveal that sex differences in the salience of anxiety persist in a second session of the TIDAL conflict test.

### In a second TIDAL session, female and male mice show expedited place avoidance

Based on the data above (see Fig. 4), we further explored whether mice exhibited signs of learning in a second TIDAL session. Therefore, data from the two-session TIDAL test in Fig. 4 were re-graphed to better visualize female and male mouse capacity for learning (**Fig. S4**). If mice exhibit learning, this would be interesting and would help inform the design of future TIDAL studies.

As mentioned, the dark-light test revealed that male (but not female) mice showed reduced dark plate preference in Session 2. As temperatures climbed, females in Session 2 showed reduced preference for the dark plate compared to the same mice completing the task in Session 1 (Females: two-way RM ANOVA with Bonferroni, session x temperature interaction, *F*_6,24_=3.18, *p* < 0.05, Session 1 vs. 2 between 42-43°C). Similarly, males in Session 2 as compared to Session 1 decreased preference for the dark plate (Males: main effect of session [*p* = 0.05] and temperature [*p* < 0.001]) (**Fig. S4A,B**). Focusing on a key temperature, 42°C, female mice in Session 1 showed 79% dark plate preference, which was reduced to 41% preference in Session 2 (two-way RM ANOVA; session x temperature interaction, *p* < 0.05) (**Fig. S5C**). At 42°C, male mice in Sessions 1 and 2 showed 44% and 28% dark plate preference, respectively (no significant differences between sessions, *p* = 0.07; two-way RM ANOVA; main effect of temperature only, *p* < 0.001) (**Fig. S4D**).

Therefore, mice completing a second session of TIDAL exhibited signs of learning by increasing dark plate avoidance: male mice in Session 2 already showed place avoidance from the dark plate at isothermic temperature at test start, whereas females in Session 2 showed accelerated avoidance as the dark plate temperature increased. Mice exhibit learning in the TIDAL conflict test, which manifests differently in female and male mice. Overall, our data suggest that the TIDAL conflict test is reliable and reproducible in showing that female as compared to male mice exhibit heightened anxiety-like behavior.

### An anxiolytic, paroxetine, reduces anxiety-like behavior in the TIDAL conflict test

Finally, we used a pharmacologic compound to validate the TIDAL conflict test as a viable strategy for assessing anxiety-like behaviors in mice (Fig. 5, **Fig. S5**). We aimed to relieve anxiety-like behavior in mice using the selective serotonin reuptake inhibitor (SSRI) paroxetine; paroxetine reduces anxiety in humans (Nemeroff & Owens, 2003; Sheehan & Mao, 2003) and mice (Bentefour et al., 2015). If paroxetine-treated mice show reduced time in the dark cylinder relative to vehicle controls, this suggests that TIDAL is valid for detecting anxiety-like behavior.

**Figure 5.**
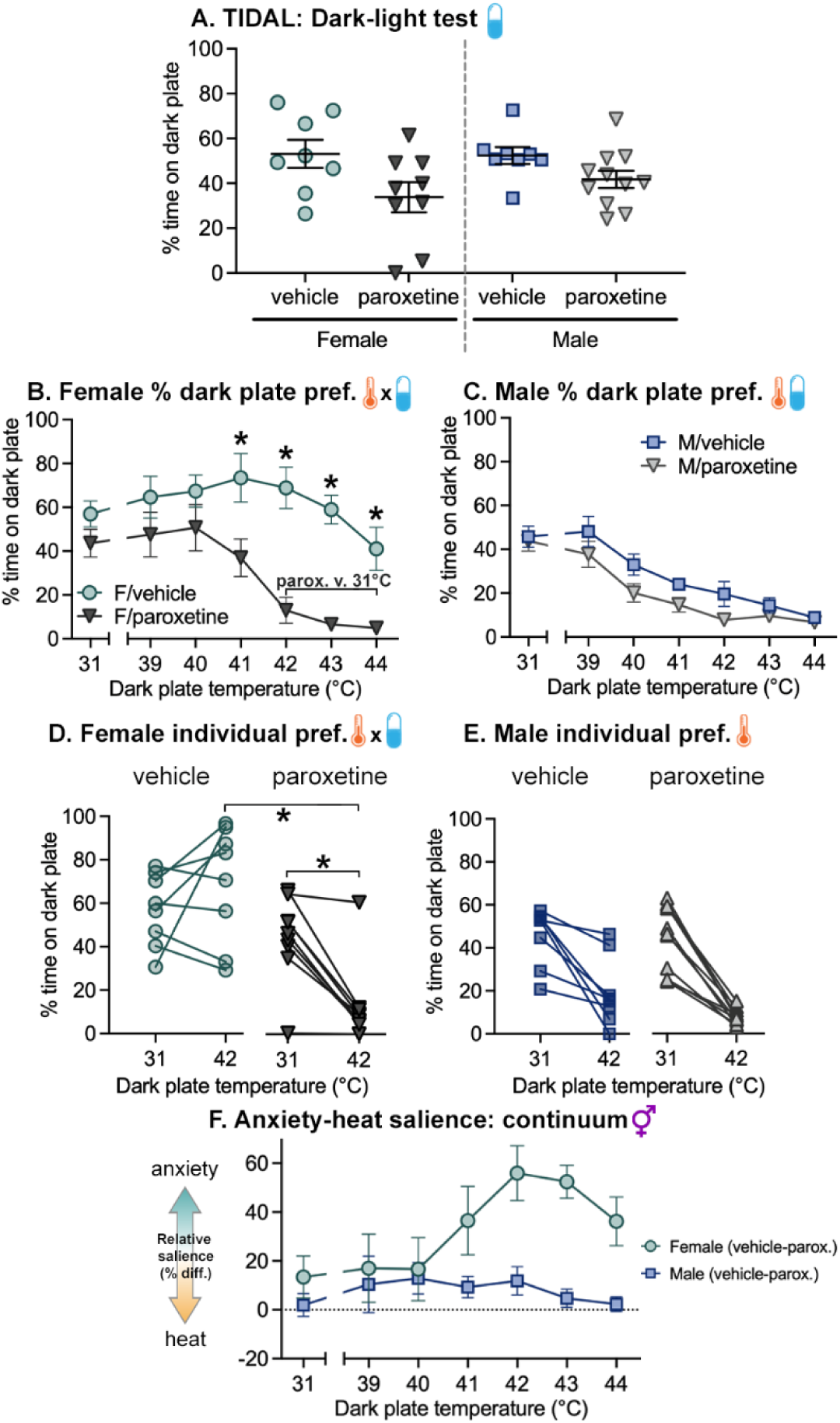
Paroxetine reduces anxiety-like behavior in female mice in the TIDAL conflict test. **A.** Mice that received paroxetine prior to TIDAL conflict testing showed reduced dark cylinder preference relative to vehicle controls with both plates at 31°C. **B.** During the TIDAL conflict test, all mice reduced time spent on the dark plate as temperatures increased. Further, female mice that received paroxetine exhibited reduced dark plate preference relative to vehicle controls from (31-44°C). **C.** Vehicle and paroxetine male mice showed similar preference for the dark plate as temperatures increased. **D-E.** Dark plate preferences of individual mice with the dark plate at 31°C and 42°C. At 42°C, vehicle females had higher dark plate preference than paroxetine females (F/vehicle: 68.9%; F/paroxetine: 12.9%). Additionally, paroxetine females showed a significant reduction in dark plate preference from 31°C to 42°C (31°C:43.6%; 42°C: 12.9%). **F.** Anxiety-heat salience continuum. Difference scores were calculated to better delineate differences between paroxetine and vehicle control groups. Subtracting paroxetine from vehicle control percent dark plate preference, female mice showed increased salience of anxiety-inducing stimulus (dark) vs. heat. F/vehicle: *n*=8 female mice, F/paroxetine *n*=9 female mice; M/vehicle: *n*=8 male mice, M/paroxetine: *n*=11 male mice. * indicates *p* < 0.05 between F/paroxetine and F/vehicle mice; “thermometer x pill” symbol indicates significant temperature x drug interaction; temperature symbol alone indicates significant main effect of temperature.

To establish whether anxiolytic treatment shifted behavior in the TIDAL conflict test, mice were administered intraperitoneal paroxetine (10 mg/kg) or saline one hour prior to TIDAL testing. In the dark-light test, mice that received paroxetine showed decreased dark plate preference compared to vehicle controls (two-way ANOVA; main effect of drug *F*_1,32_ = 8.13, *p* < 0.05) (Fig. 5A). During TIDAL conflict testing, female-paroxetine (vs. female-vehicle) mice decreased preference for the heated dark plate from 41-44°C (two-way RM ANOVA, drug x temperature interaction, *F*_6,90_=3.80, *p* <0.005, female-vehicle vs. female-paroxetine between 41-44°C, female-paroxetine 42-44°C vs. 31°C) (Fig. 5B). Male-paroxetine mice also decreased preference for the dark plate relative to vehicle controls (two-way RM ANOVA, main effect of drug (*F*_1,102_=4.62, *p* < 0.05) and temperature (*F*_6,102_=35.53, *p* < 0.005)) (Fig. 5C). In using paroxetine in the TIDAL conflict test, 42°C robustly uncovered differences in anxiety-like behavior between treatment groups: in mice at 42°C, dark plate preference was 69% for female-vehicle and 13% for female-paroxetine mice, and male dark plate preference was 20% for male-vehicle and 8% for male-paroxetine mice, respectively (Fig. 5D**,E**). Female-vehicle mice preferred the dark plate significantly more than female-paroxetine mice at 42°C. Additionally, whereas female-vehicle mice showed similar dark plate preference at 31°C (57%) and 42°C (69%), female-paroxetine mice reduced dark plate preference from 44% at baseline 31°C to 8% at 42°C (two-way RM ANOVA, drug x temperature interaction, *F*_1,15_=15.09, *p* < 0.005, female-vehicle vs. female-paroxetine at 42°C, paroxetine female 31°C vs. 42°C). All male mice regardless of drug treatment exhibited decreased dark plate preference from 31°C to 44°C (main effect of temperature, *F*_1,17_=69.74, *p* < 0.005).

To further examine the relative salience of anxiety vs. heat hypersensitivity in vehicle-and paroxetine-treated mice, we calculated difference scores (Fig. 5E). Subtracting vehicle minus paroxetine scores, these difference scores revealed a robust anxiolytic effect of paroxetine especially in female mice (two-way ANOVA, main effect of sex, *F*_1,6_=13. 48, p < 0.05).

Averaging from 42-44°C, female-paroxetine mice showed 48% reduced dark plate preference compared to female-vehicle mice. From 39-42°C, male-paroxetine mice had 11% decreased dark plate preference compared to male-vehicle mice. These results reveal that paroxetine reduces dark plate preference as temperatures rise in the TIDAL conflict test, suggesting that the TIDAL conflict test for mice is valid for assessing anxiety-like behavior.

## Discussion

This study explored anxiety-like behavior in mice using the TIDAL conflict test, a novel assay integrating a conflicting dark-light dilemma with incremental increases on a heating plate. The TIDAL conflict test unmasked robust and reproducible sex differences in anxiety: female as compared to male mice maintained prolonged dark plate preference under increasingly aversive hyperthermic conditions, suggesting increased anxiety-like behaviors. When mice completed a second session of TIDAL, females showed increased anxiety-like symptoms in the dark-light test (compared to the dark-light test prior to the first session). Further, mice of both sexes in a second session had accelerated place avoidance of the heating dark plate, implying that they had learned from prior exposure to the test; thus, to avoid confounds of learning, distinct cohorts should be used for testing manipulations or timecourses. Anxiety-related shifts in TIDAL behavior were not simply a preference for the heated plate, since mice exhibited prolonged preference for the heated plate in the TIDAL conflict test (heated plate dark; isothermic plate illuminated) compared to the TPP test (both sides illuminated). Finally, we validated the TIDAL conflict test using an anxiety-relieving drug, paroxetine, which decreased mouse preference for the dark-heating plate. Therefore, compared to one-dimensional, commonly used tests of anxiety behaviors, our newly established TIDAL conflict test reveals tangible differences in anxiety-like symptoms in mice due to sex.

Differences in human anxiety are not consistently replicated in rodent models (Donner & Lowry, 2013; Scholl et al., 2019). Most tests for anxiety-like behavior were developed >20 years ago and were validated in male rodents only (Börchers et al., 2022; Donner & Lowry, 2013). In the elevated plus maze, female mice show decreased anxiety-like behavior (defined as increased time in open arms and open arm entries) when compared to male mice (Rodgers & Cole, 1993; Võikar et al., 2001). However, in the light-dark test, mouse anxiety-like behavior is variable between sexes across different mouse strains (Võikar et al., 2001). Accordingly, we found that the light-dark test – completed in the first five minutes in the apparatus with both plates at 31°C – showed little or no difference between sexes; sex differences in anxiety-like behavior were only unmasked as the anxiety-driving stimulus was placed in conflict with an aversive temperature stimulus. This corroborates previous studies described above and underscores a need to develop more refined tests that model anxiety in mice.

Here, we sought to develop a conflict test in mice that better uncovered differences in anxiety-like behavior. We placed an anxiety-relevant dilemma (light vs. dark) in conflict with increasing temperature (on the dark side only). Our optimized TIDAL conflict test incorporates dark plate temperatures initially at 31°C (10 min), then incrementally increasing temperatures of 39°C-44°C (5 min each). Our optimized temperature range is notable, because the heat-activated ion channel involved in sensing low-level noxious heat, TRPV1, is activated at 43°C (Willis, 2009). The TIDAL conflict test revealed robust, reproducible increases in anxiety-like behavior in females vs. males. Female rodents exhibit increased pain symptoms on reflexive tests (Hargreaves test or hot plate) (Gaudet et al., 2017; Gioiosa et al., 2008; Mogil, 2020) but also prefer warmer ambient temperatures (Kaikaew et al., 2017) – these sex differences in temperature preferences could influence TIDAL outcomes. Our control conditions confirmed that the observed sex differences were not simply due to enhanced preference for the heated plate for females; females on TIDAL displayed stronger preference for the heated (also dark) plate compared to females on TPP (with the heated plated lighted). Therefore, the TIDAL conflict test is a novel approach in mice for uncovering sex differences in anxiety-like behavior.

Mice of both sexes exhibited learning on the TIDAL conflict test. Mice were tested in two sessions separated by two weeks using identical TIDAL conflict testing protocols. Learning was apparent in the second session in the initial dark-light test: female mice in the second dark-light test newly showed increased dark plate preference compared to males. In TIDAL Sessions 1 and 2, female as compared to male mice showed increased preference for the heat shift-dark plate from 39°C throughout the remainder of the test. Further, in the second TIDAL session, both females and males expedited exit from the heat shift-dark plate, implying that they anticipated the increasing temperatures and proactively avoided this side. These results extend our previous findings that females as compared to male exhibit robust anxiety-like behavior in the TIDAL conflict assay – in particular, females showed heightened anxiety-like behavior in the Session 2 dark-light test. Further, our results show that mice learn the test, and that sex differences in the salience of anxiety persist through repeated TIDAL sessions. Similarly, rodent learning occurs in other tests of anxiety-like behavior (Bailey & Crawley, 2009; File, 1993, 2001; Roesler et al., 1999). The fact that rodents exhibit learning on the TIDAL conflict test is important, because this suggests that behavioral timecourses or tests using different treatments must account for this learning effect or experiments must be completed on distinct cohorts.

Our TIDAL conflict test uncovered sex differences in mice that align well with sex differences in anxiety observed in humans. In the human population, the lifetime prevalence of anxiety disorders is up to 60% higher in women (versus men) (Kessler et al., 2005; McLean & Anderson, 2009). Symptom progression, treatment response, and average age of onset are also affected by gender (Pigott, 2003). In our TIDAL conflict test, female mice showed increased anxiety-like behavior; however, the basic light-dark test – which is frequently used to assay anxiety-like behavior – failed to reliably detect sex differences. This suggests that a more complex test is required to identify behavioral differences in mice of different sexes, which is supported by the differences in clinical presentation observed across genders in humans.

Anxiety disorders often present comorbidly with other health conditions such as depression, hypertension, epilepsy, chronic pain, and neurotrauma (Dickerson et al., 2021; Hingray et al., 2019; Johnson, 2019; Tiller, 2013). We did not assess estrous cycle, which is accordance with guidelines for studying sex differences in rodents (Shansky, 2019; Shansky & Murphy, 2021) since gonadal hormones were not the main question studied here. Future studies could further explore the role of hormones in anxiety-like behavior using the TIDAL conflict test. Overall, it is important to identify mechanistic differences across genders that underlie susceptibility to anxiety and comorbid conditions.

We used the TIDAL conflict test previously to evaluate anxiety-like behavior of mice with spinal cord injury, which revealed that mice with spinal cord injury exhibited increased anxiety-like behavior relative to sham surgery controls (Lee et al., 2023). Here, we sought to further validate the TIDAL conflict test using known pharmacologic agents that ameliorate anxiety-like behavior. Initially, we attempted validation using diazepam, a commonly used benzodiazepine that modulates GABA influx (Campo-Soria et al., 2006; Lavoie & Twyman, 1996; Masiulis et al., 2019). In C57BL/6J mice, however, diazepam can cause strong sedative effects that mask potential anxiolytic effects (Pádua-Reis et al., 2021). In our pilot studies delivering diazepam prior to the TIDAL conflict test, all diazepam-treated mice substantially decreased overall exploration and reduced time spent in the dark cylinder (data not shown). Therefore, acute diazepam induces hypolocomotion that impedes ability to assess anxiety-like behavior in our place preference tests.

Next, we tested the SSRI paroxetine. SSRIs can treat clinical anxiety without producing severe locomotor side effects (Jakubovski et al., 2019; Sheehan & Mao, 2003). Following acute paroxetine administration, mice exhibited reduced overall exploration and time spent in the dark cylinder during the TIDAL conflict test. This result was more pronounced in female mice, suggesting underlying sex differences play a role in anxiety-like behavioral state. It is important to note that in humans, paroxetine treatment is typically a long-term intervention, often eliciting noticeable behavioral changes within 1-3 months (Perna et al., 2016). In contrast, here, mice received a single dose of paroxetine one hour prior to TIDAL conflict testing, which replicates acute anxiolytic effects in mice observed previously (Pádua-Reis et al., 2021). Future studies could assess anxiolytic effectiveness of paroxetine as a long-term intervention in male and female mice. Overall, these results using a non-sedative anxiolytic confirm the validity and reliability of the TIDAL conflict test for assessing anxiety-like behavior.

As mentioned, the TIDAL conflict test more effectively unmasks clinically relevant sex differences in anxiety-like behavior than other tests, and this is reflected in effect sizes. To contextualize the observed sex difference in TIDAL, we calculated the standardized mean difference score (d) (Cohen, 2013) based on the 50% threshold temperature values from Fig. 3B. The resulting value of d = 2.68, with females more strongly preferring the dark, heating chamber compared to males, suggests a relatively “large” effect size (Sullivan & Feinn, 2012). In contrast, in the open field test, little-to-no sex differences are observed in percent time in the center zone (Fritz et al., 2017; Vošlajerová Bímová et al., 2016). In the elevated plus maze, an examination of reported sex differences reveals the following: Rogers & Cole (1993) demonstrate that females spend significantly less time in the closed arms relative to males, yielding a difference score of 0.798, and more time in the open arms (d = 0.53), though the difference is non-significant; Hendershott et al. (2016) report that females exhibited an increased preference for the open arms with a maximum difference score of 0.868; and Painsipp et al. (2007) similarly note females’ heightened preference for the open arms, with a difference score of d = 1.54. The significant sex difference reported in these studies fall near or within the “large” range (≥ 0.8) (Cohen, 2013; Sullivan & Feinn, 2012); however, these results are in the opposite direction of our finding that females exhibit more anxiety-like behavior than males. Our larger effect size implies less overlap between females in males in the TIDAL conflict test compared to the elevated plus maze, and our TIDAL sex differences are in a direction that parallels human gender differences in anxiety prevalence. Ultimately, this suggests that the sex differences in anxiety-like behavior detected by TIDAL are pronounced and clinically relevant, highlighting unique aspects of anxiety-related responses in this test compared to commonly used tests.

Our results highlight the importance of conflict tests in uncovering anxiety-like behavioral differences in mice of different groups. Conflict tests produce differing motivational states through the introduction of approach-avoidance situations. These tests offer an unconditioned approach to observing anxiety-like behavior, resulting in high ethological validity (Campos et al., 2013). Previous conflict tests have assayed anxiety-like behavior, including the Vogel test (Basso et al., 2011; Johnston & File, 1991; Vogel et al., 1971), the four-plate assay (Boissier et al., 1968), and the defensive burying test (Fucich & Morilak, 2018; Pinel & Treit, 1978). There are key considerations and limitations related to these existing conflict tests (Lapiz-Bluhm et al., 2008): (1) each test uses shock as a punishment, which may cause pain and assess fear (rather than anxiety); (2) several tests have not been rigorously validated in female vs. male rodents (four-plate test) or do not show clearly interpretable sex differences (defensive burying; Arakawa, 2007; Castillo et al., 2022); (3) the four-plate test elicits many false positives (File, 2001); and (4) the Vogel and probe burying tests were optimized for study in rats, rather than mice. Thus, benefits of the TIDAL conflict test include that it does not require deprivation or training or induce pain; it is validated for use in mice; and the test mouse is free to explore the aversive chamber or not, thereby limiting stressful effects of the test.

## Future directions and conclusions

Here, we developed and validated a new assay – the thermal increments dark-light (TIDAL) conflict test – that exposes in mice previously underappreciated differences in anxiety-like behavior. Future studies could use this test to explore neural circuitry related to anxiety or avoidance behaviors (Bangasser & Cuarenta, 2021); e.g., to test whether manipulating specific neural pathways alters anxiety-like behavior in TIDAL. Further, TIDAL experiments could explore anxiety-like behavior in other contexts, including stress, early-life adversity, injury, or sickness/neuroinflammation (Bolton et al., 2018; Bourke et al., 2012; Fonken et al., 2013, 2018; Grace et al., 2021).

In summary, we explored anxiety-like behavior in mice using the novel heat-light TIDAL conflict test. Our data reveal that the TIDAL conflict test reliably unmasks amplified anxiety symptoms in mice that are female compared to males; the test was validated using an anxiety-relieving drug, which reduced mouse preference for the dark-heating plate. Incrementally increasing the magnitude of one of the conflicting factors (here, heat), while maintaining constant the other factor (dark vs. light), enabled deciphering robust differences that might have been overlooked if only a single temperature was used. More broadly, these results suggest that rodent studies should incorporate conflicting stimuli to illuminate potential differences in anxiety-like behavior. Therefore, future preclinical studies should prioritize assays that detect behavioral differences not apparent in commonly used anxiety-like behavioral assays to identify circuits and therapies that benefit health outcomes, emotive state, and well-being.

## Acknowledgements

We thank the Animal Resources Center (ARC) husbandry staff at the Health Discovery Building for excellent animal care. Partial support was provided by University of Texas at Austin start-up funds (ADG), the Wings for Life Foundation (ADG), and Mission Connect, a program of the TIRR Foundation (ADG). Research reported in this publication was supported by the National Institute Of Neurological Disorders And Stroke of the National Institutes of Health under Award Number R01NS131806 (ADG), and by National Institutes of Health Awards R01AG062716 (LKF) and R01AG078758 (LKF). The content is solely the responsibility of the authors and does not necessarily represent the official views of the National Institutes of Health.

## Author contributions

SEL, SHP, LKF, & ADG designed experiments. SEL & SHP performed experiments. SEL, JCA, SHP, & ADG analyzed data. SEL, JCA, SHP, LKF, & ADG wrote and edited the manuscript.

## References

1. Altemus, M., Sarvaiya, N., & Neill Epperson, C. (2014). Sex differences in anxiety and depression clinical perspectives. Frontiers in Neuroendocrinology, 35(3), 320–330. 10.1016/j.yfrne.2014.05.004

2. An, X.-L., Zou, J.-X., Wu, R.-Y., Yang, Y., Tai, F.-D., Zeng, S.-Y., Jia, R., Zhang, X., Liu, E.-Q., & Broders, H. (2011). Strain and sex differences in anxiety-like and social behaviors in C57BL/6J and BALB/cJ mice. Experimental Animals, 60(2), 111–123. 10.1538/expanim.60.111

3. Arakawa, H. (2007). Ontogeny of sex differences in defensive burying behavior in rats: Effect of social isolation. Aggressive Behavior, 33(1), 38–47. 10.1002/ab.20165

4. Bailey, K. R., & Crawley, J. N. (2009). Anxiety-Related Behaviors in Mice. In J. J. Buccafusco (Ed.), Methods of Behavior Analysis in Neuroscience (2nd ed.). CRC Press/Taylor & Francis. http://www.ncbi.nlm.nih.gov/books/NBK5221/

5. Bangasser, D. A., & Cuarenta, A. (2021). Sex differences in anxiety and depression: Circuits and mechanisms. Nature Reviews. Neuroscience, 22(11), 674–684. 10.1038/s41583-021-00513-0

6. Bartley, E. J., & Fillingim, R. B. (2013). Sex differences in pain: A brief review of clinical and experimental findings. British Journal of Anaesthesia, 111(1), 52–58. 10.1093/bja/aet127

7. Basso, A. M., Gallagher, K. B., Mikusa, J. P., & Rueter, L. E. (2011). Vogel conflict test: Sex differences and pharmacological validation of the model. Behavioural Brain Research, 218(1), 174–183. 10.1016/j.bbr.2010.11.041

8. Baxter, A. J., Scott, K. M., Vos, T., & Whiteford, H. A. (2013). Global prevalence of anxiety disorders: A systematic review and meta-regression. Psychological Medicine, 43(5), 897–910. 10.1017/S003329171200147X

9. Bekhbat, M., & Neigh, G. N. (2018). Sex differences in the neuro-immune consequences of stress: Focus on depression and anxiety. Brain, Behavior, and Immunity, 67, 1–12. 10.1016/j.bbi.2017.02.006

10. Bentefour, Y., Bennis, M., Garcia, R., & M’hamed, S. B. (2015). Effects of paroxetine on PTSD-like symptoms in mice. Psychopharmacology, 232(13), 2303–2312. 10.1007/s00213-014-3861-2

11. Boissier, J. R., Simon, P., & Aron, C. (1968). A new method for rapid screening of minor tranquillizers in mice. European Journal of Pharmacology, 4(2), 145–151. 10.1016/0014-2999(68)90170-2

12. Bolton, J. L., Molet, J., Regev, L., Chen, Y., Rismanchi, N., Haddad, E., Yang, D. Z., Obenaus, A., & Baram, T. Z. (2018). Anhedonia Following Early-Life Adversity Involves Aberrant Interaction of Reward and Anxiety Circuits and Is Reversed by Partial Silencing of Amygdala Corticotropin-Releasing Hormone Gene. Biological Psychiatry, 83(2), 137–147. 10.1016/j.biopsych.2017.08.023

13. Börchers, S., Krieger, J.-P., Asker, M., Maric, I., & Skibicka, K. P. (2022). Commonly-used rodent tests of anxiety-like behavior lack predictive validity for human sex differences. Psychoneuroendocrinology, 141, 105733. 10.1016/j.psyneuen.2022.105733

14. Bourke, C. H., Harrell, C. S., & Neigh, G. N. (2012). Stress-induced sex differences: Adaptations mediated by the glucocorticoid receptor. Hormones and Behavior, 62(3), 210–218. 10.1016/j.yhbeh.2012.02.024

15. Bragdon, E. E., Light, K. C., Costello, N. L., Sigurdsson, A., Bunting, S., Bhalang, K., & Maixner, W. (2002). Group differences in pain modulation: Pain-free women compared to pain-free men and to women with TMD. Pain, 96(3), 227–237. 10.1016/S0304-3959(01)00451-1

16. Campos, A. C., Fogaça, M. V., Aguiar, D. C., & Guimarães, F. S. (2013). Animal models of anxiety disorders and stress. Brazilian Journal of Psychiatry, 35, S101–S111. 10.1590/1516-4446-2013-1139

17. Campo-Soria, C., Chang, Y., & Weiss, D. S. (2006). Mechanism of action of benzodiazepines on GABAA receptors. British Journal of Pharmacology, 148(7), 984–990. 10.1038/sj.bjp.0706796

18. Castillo, L. Y., Ríos-Carrillo, J., González-Orozco, J. C., Camacho-Arroyo, I., Morin, J.-P., Zepeda, R. C., & Roldán-Roldán, G. (2022). Juvenile Exposure to BPA Alters the Estrous Cycle and Differentially Increases Anxiety-like Behavior and Brain Gene Expression in Adult Male and Female Rats. Toxics, 10(9). 10.3390/toxics10090513

19. Cohen, J. (2013). *Statistical Power Analysis for the Behavioral Sciences* (2nd ed.). Routledge. 10.4324/9780203771587

20. Dickerson, M. R., Murphy, S. F., Urban, M. J., White, Z., & VandeVord, P. J. (2021). Chronic Anxiety- and Depression-Like Behaviors Are Associated With Glial-Driven Pathology Following Repeated Blast Induced Neurotrauma. Frontiers in Behavioral Neuroscience, 15, 787475. 10.3389/fnbeh.2021.787475

21. Dieleman, J. L., Baral, R., Birger, M., Bui, A. L., Bulchis, A., Chapin, A., Hamavid, H., Horst, C., Johnson, E. K., Joseph, J., Lavado, R., Lomsadze, L., Reynolds, A., Squires, E., Campbell, M., DeCenso, B., Dicker, D., Flaxman, A. D., Gabert, R., … Murray, C. J. L. (2016). US Spending on Personal Health Care and Public Health, 1996-2013. JAMA, 316(24), 2627–2646. 10.1001/jama.2016.16885

22. Donner, N. C., & Lowry, C. A. (2013). Sex differences in anxiety and emotional behavior. Pflügers Archiv - European Journal of Physiology, 465(5), 601–626. 10.1007/s00424-013-1271-7

23. Feine, J. S., Bushnell, C. M., Miron, D., & Duncan, G. H. (1991). Sex differences in the perception of noxious heat stimuli. Pain, 44(3), 255–262. 10.1016/0304-3959(91)90094-E

24. File, S. E. (1993). The interplay of learning and anxiety in the elevated plus-maze. Behavioural Brain Research, 58(1–2), 199–202. 10.1016/0166-4328(93)90103-w

25. File, S. E. (2001). Factors controlling measures of anxiety and responses to novelty in the mouse. Behavioural Brain Research, 125(1–2), 151–157. 10.1016/s0166-4328(01)00292-3

26. Fonken, L. K., Frank, M. G., Gaudet, A. D., D’Angelo, H. M., Daut, R. A., Hampson, E. C., Ayala, M. T., Watkins, L. R., & Maier, S. F. (2018). Neuroinflammatory priming to stress is differentially regulated in male and female rats. Brain, Behavior, and Immunity, 70, 257–267. 10.1016/j.bbi.2018.03.005

27. Fonken, L. K., Weil, Z. M., & Nelson, R. J. (2013). Mice exposed to dim light at night exaggerate inflammatory responses to lipopolysaccharide. Brain, Behavior, and Immunity, 34, 159–163. 10.1016/j.bbi.2013.08.011

28. Fritz, A.-K., Amrein, I., & Wolfer, D. P. (2017). Similar reliability and equivalent performance of female and male mice in the open field and water-maze place navigation task. American Journal of Medical Genetics. Part C, Seminars in Medical Genetics, 175(3), 380–391. 10.1002/ajmg.c.31565

29. Fucich, E. A., & Morilak, D. A. (2018). Shock-probe Defensive Burying Test to Measure Active versus Passive Coping Style in Response to an Aversive Stimulus in Rats. Bio-Protocol, 8(17), e2998. 10.21769/BioProtoc.2998

30. Gaudet, A. D., Ayala, M. T., Schleicher, W. E., Smith, E. J., Bateman, E. M., Maier, S. F., & Watkins, L. R. (2017). Exploring acute-to-chronic neuropathic pain in rats after contusion spinal cord injury. Experimental Neurology, 295, 46–54. 10.1016/j.expneurol.2017.05.011

31. Gioiosa, L., Chen, X., Watkins, R., Klanfer, N., Bryant, C. D., Evans, C. J., & Arnold, A. P. (2008). Sex chromosome complement affects nociception in tests of acute and chronic exposure to morphine in mice. Hormones and Behavior, 53(1), 124–130. 10.1016/j.yhbeh.2007.09.003

32. Grace, P. M., Tawfik, V. L., Svensson, C. I., Burton, M. D., Loggia, M. L., & Hutchinson, M. R. (2021). The Neuroimmunology of Chronic Pain: From Rodents to Humans. The Journal of Neuroscience: The Official Journal of the Society for Neuroscience, 41(5), 855–865. 10.1523/JNEUROSCI.1650-20.2020

33. Hendershott, T. R., Cronin, M. E., Langella, S., McGuinness, P. S., & Basu, A. C. (2016). Effects of environmental enrichment on anxiety-like behavior, sociability, sensory gating, and spatial learning in male and female C57BL/6J mice. Behavioural Brain Research, 314, 215–225. 10.1016/j.bbr.2016.08.004

34. Hingray, C., McGonigal, A., Kotwas, I., & Micoulaud-Franchi, J.-A. (2019). The Relationship Between Epilepsy and Anxiety Disorders. Current Psychiatry Reports, 21(6), 40. 10.1007/s11920-019-1029-9

35. Jakubovski, E., Johnson, J. A., Nasir, M., Müller-Vahl, K., & Bloch, M. H. (2019). Systematic review and meta-analysis: Dose-response curve of SSRIs and SNRIs in anxiety disorders. Depression and Anxiety, 36(3), 198–212. 10.1002/da.22854

36. Johnson, H. M. (2019). Anxiety and Hypertension: Is There a Link? A Literature Review of the Comorbidity Relationship Between Anxiety and Hypertension. Current Hypertension Reports, 21(9), 66. 10.1007/s11906-019-0972-5

37. Johnston, A. L., & File, S. E. (1991). Sex differences in animal tests of anxiety. Physiology & Behavior, 49(2), 245–250. 10.1016/0031-9384(91)90039-q

38. Kaikaew, K., Steenbergen, J., Themmen, A. P. N., Visser, J. A., & Grefhorst, A. (2017). Sex difference in thermal preference of adult mice does not depend on presence of the gonads. Biology of Sex Differences, 8(1), 24. 10.1186/s13293-017-0145-7

39. Kessler, R. C., Chiu, W. T., Demler, O., Merikangas, K. R., & Walters, E. E. (2005). Prevalence, severity, and comorbidity of 12-month DSM-IV disorders in the National Comorbidity Survey Replication. Archives of General Psychiatry, 62(6), 617–627. 10.1001/archpsyc.62.6.617

40. Knight, P., Chellian, R., Wilson, R., Behnood-Rod, A., Panunzio, S., & Bruijnzeel, A. W. (2021). Sex differences in the elevated plus-maze test and large open field test in adult Wistar rats. Pharmacology, Biochemistry, and Behavior, 204, 173168. 10.1016/j.pbb.2021.173168

41. Lapiz-Bluhm, M. D. S., Bondi, C. O., Doyen, J., Rodriguez, G. A., Bédard-Arana, T., & Morilak, D. A. (2008). Behavioural assays to model cognitive and affective dimensions of depression and anxiety in rats. Journal of Neuroendocrinology, 20(10), 1115–1137. 10.1111/j.1365-2826.2008.01772.x

42. Lavoie, A. M., & Twyman, R. E. (1996). Direct evidence for diazepam modulation of GABAA receptor microscopic affinity. Neuropharmacology, 35(9–10), 1383–1392. 10.1016/s0028-3908(96)00077-9

43. Lee, S. E., Greenough, E. K., Fonken, L. K., & Gaudet, A. D. (2023). Spinal cord injury in mice amplifies anxiety: A novel light-heat conflict test exposes increased salience of anxiety over heat. BioRxiv, 2023.01.13.523970. 10.1101/2023.01.13.523970

44. Masiulis, S., Desai, R., Uchański, T., Serna Martin, I., Laverty, D., Karia, D., Malinauskas, T., Zivanov, J., Pardon, E., Kotecha, A., Steyaert, J., Miller, K. W., & Aricescu, A. R. (2019). GABAA receptor signalling mechanisms revealed by structural pharmacology. Nature, 565(7740), 454–459. 10.1038/s41586-018-0832-5

45. McLean, C. P., & Anderson, E. R. (2009). Brave men and timid women? A review of the gender differences in fear and anxiety. Clinical Psychology Review, 29(6), 496–505. 10.1016/j.cpr.2009.05.003

46. McLean, C. P., Asnaani, A., Litz, B. T., & Hofmann, S. G. (2011). Gender differences in anxiety disorders: Prevalence, course of illness, comorbidity and burden of illness. Journal of Psychiatric Research, 45(8), 1027–1035. 10.1016/j.jpsychires.2011.03.006

47. Mogil, J. S. (2020). Qualitative sex differences in pain processing: Emerging evidence of a biased literature. Nature Reviews. Neuroscience, 21(7), 353–365. 10.1038/s41583-020-0310-6

48. Nemeroff, C. B., & Owens, M. J. (2003). Neuropharmacology of paroxetine. Psychopharmacology Bulletin, 37 *Suppl 1*, 8–18.

49. Pádua-Reis, M., Nôga, D. A., Tort, A. B. L., & Blunder, M. (2021). Diazepam causes sedative rather than anxiolytic effects in C57BL/6J mice. Scientific Reports, 11(1), 9335. 10.1038/s41598-021-88599-5

50. Painsipp, E., Wultsch, T., Shahbazian, A., Edelsbrunner, M., Kreissl, M. C., Schirbel, A., Bock, E., Pabst, M. A., Thoeringer, C. K., Huber, H. P., & Holzer, P. (2007). Experimental gastritis in mice enhances anxiety in a gender-related manner. Neuroscience, 150(3), 522–536. 10.1016/j.neuroscience.2007.09.024

51. Perna, G., Alciati, A., Riva, A., Micieli, W., & Caldirola, D. (2016). Long-Term Pharmacological Treatments of Anxiety Disorders: An Updated Systematic Review. Current Psychiatry Reports, 18(3), 23. 10.1007/s11920-016-0668-3

52. Pigott, T. A. (2003). Anxiety disorders in women. Psychiatric Clinics of North America, 26(3), 621–672. 10.1016/S0193-953X(03)00040-6

53. Pinel, J. P., & Treit, D. (1978). Burying as a defensive response in rats. Journal of Comparative and Physiological Psychology, 92(4), 708–712. 10.1037/h0077494

54. Reddan, M. C., Young, H., Falkner, J., López-Solà, M., & Wager, T. D. (2020). Touch and social support influence interpersonal synchrony and pain. Social Cognitive and Affective Neuroscience, 15(10), 1064–1075. 10.1093/scan/nsaa048

55. Rhudy, J. L., & Meagher, M. W. (2001). Noise stress and human pain thresholds: Divergent effects in men and women. The Journal of Pain, 2(1), 57–64. 10.1054/jpai.2000.19947

56. Rodgers, R. J., & Cole, J. C. (1993). Influence of social isolation, gender, strain, and prior novelty on plus-maze behaviour in mice. Physiology & Behavior, 54(4), 729–736. 10.1016/0031-9384(93)90084-s

57. Roesler, R., Walz, R., Quevedo, J., de-Paris, F., Zanata, S. M., Graner, E., Izquierdo, I., Martins, V. R., & Brentani, R. R. (1999). Normal inhibitory avoidance learning and anxiety, but increased locomotor activity in mice devoid of PrP(C). Brain Research. Molecular Brain Research, 71(2), 349–353. 10.1016/s0169-328x(99)00193-x

58. Scholl, J. L., Afzal, A., Fox, L. C., Watt, M. J., & Forster, G. L. (2019). Sex differences in anxiety-like behaviors in rats. Physiology & Behavior, 211, 112670. 10.1016/j.physbeh.2019.112670

59. Shansky, R. M. (2019). Are hormones a “female problem” for animal research? Science (New York, N.Y.), 364(6443), 825–826. 10.1126/science.aaw7570

60. Shansky, R. M., & Murphy, A. Z. (2021). Considering sex as a biological variable will require a global shift in science culture. Nature Neuroscience, 24(4), 457–464. 10.1038/s41593-021-00806-8

61. Sheehan, D. V., & Mao, C. G. (2003). Paroxetine treatment of generalized anxiety disorder. Psychopharmacology Bulletin, 37 *Suppl 1*, 64–75.

62. Somers, J. M., Goldner, E. M., Waraich, P., & Hsu, L. (2006). Prevalence and incidence studies of anxiety disorders: A systematic review of the literature. Canadian Journal of Psychiatry. Revue Canadienne De Psychiatrie, 51(2), 100–113. 10.1177/070674370605100206

63. Sullivan, G. M., & Feinn, R. (2012). Using Effect Size—Or Why the P Value Is Not Enough. Journal of Graduate Medical Education, 4(3), 279–282. 10.4300/JGME-D-12-00156.1

64. Tiller, J. W. G. (2013). Depression and anxiety. The Medical Journal of Australia, 199(S6), S28–31. 10.5694/mja12.10628

65. Trautmann, S., Rehm, J., & Wittchen, H. (2016). The economic costs of mental disorders. EMBO Reports, 17(9), 1245–1249. 10.15252/embr.201642951

66. Vogel, J. R., Beer, B., & Clody, D. E. (1971). A simple and reliable conflict procedure for testing anti-anxiety agents. Psychopharmacologia, 21(1), 1–7. 10.1007/BF00403989

67. Võikar, V., Kõks, S., Vasar, E., & Rauvala, H. (2001). Strain and gender differences in the behavior of mouse lines commonly used in transgenic studies. Physiology & Behavior, 72(1), 271–281. 10.1016/S0031-9384(00)00405-4

68. Vošlajerová Bímová, B., Mikula, O., Macholán, M., Janotová, K., & Hiadlovská, Z. (2016). Female House Mice do not Differ in Their Exploratory Behaviour from Males. Ethology, 122(4), 298–307. 10.1111/eth.12462

69. Willis, W. D. (2009). The role of TRPV1 receptors in pain evoked by noxious thermal and chemical stimuli. Experimental Brain Research, 196(1), 5–11. 10.1007/s00221-009-1760-2

