## Supplementary material for "Anxiety-like behaviors in mice unmasked: Revealing sex differences in anxiety using a novel light-heat conflict test": Fig. S

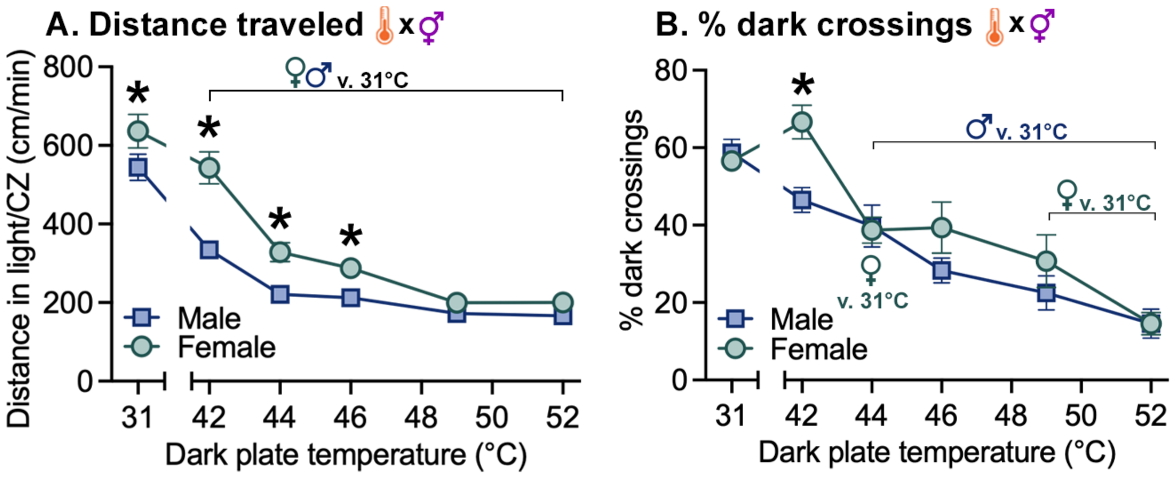


**Figure S1.** TIDAL experiment 1: as the temperature on the dark plate increased, female and male mice reduced distance moved and reduced percent dark crossings. **A.** Distance moved over temperatures/time; mice reduced distance moved as the dark plate temperature increased (and as they were on the plate for more time). Females showed increased distance moved, particularly in the lower range of temperatures. **B.** With increasing dark plate temperature, female and male mice showed reduced dark plate crossings (percent of all crossings). Females showed higher percent dark crossings at 42°C. $n$=13 female, $n$=15 male mice * indicates $p < 0.05$ between female and male mice; “female/male symbol v. 31°C” indicates within-sex difference vs. baseline 31°C.

* indicates $p < 0.05$ between female and male mice; “gender x TEST” symbol indicates significant sex x test interaction; thermometer or TEST symbols alone indicate significant main effects of temperature and TPP/TIDAL, respectively.

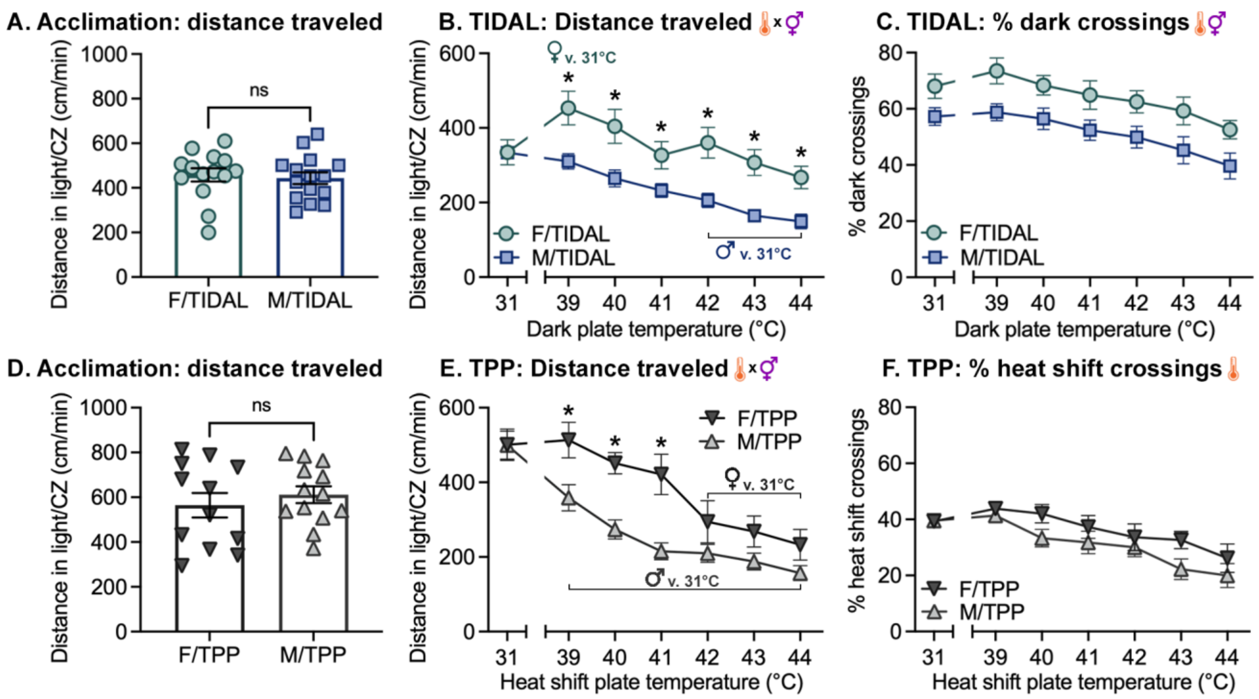


**Figure S2.** In TIDAL, mice of both sexes move less distance over time and females compared to males show increased movement and crossing into the heating-dark plate. **A,D.** Females and males travelled similar distances in the illuminated area in the acclimation period (dark-light test). **B.** In TIDAL, mice of both sexes reduced movement with time; females traveled more per minute with the heating-dark plate. **C.** In TIDAL, females crossed more frequently into the dark-heated plate. **E.** In TPP, females moved further than males, and both decreased movement as the test proceeded. **F.** In TPP, females and males exhibited a similar decrease in percent crossings onto the dark plate as the temperatures increased.

* indicates $p < 0.05$ between female and male mice; "thermometer x gender" symbol indicates significant temperature x sex interaction; thermometer or gender symbols alone indicate significant main effects of temperature and sex, respectively; "female/male symbol v. 31°C" indicates within-sex difference vs. baseline 31°C.

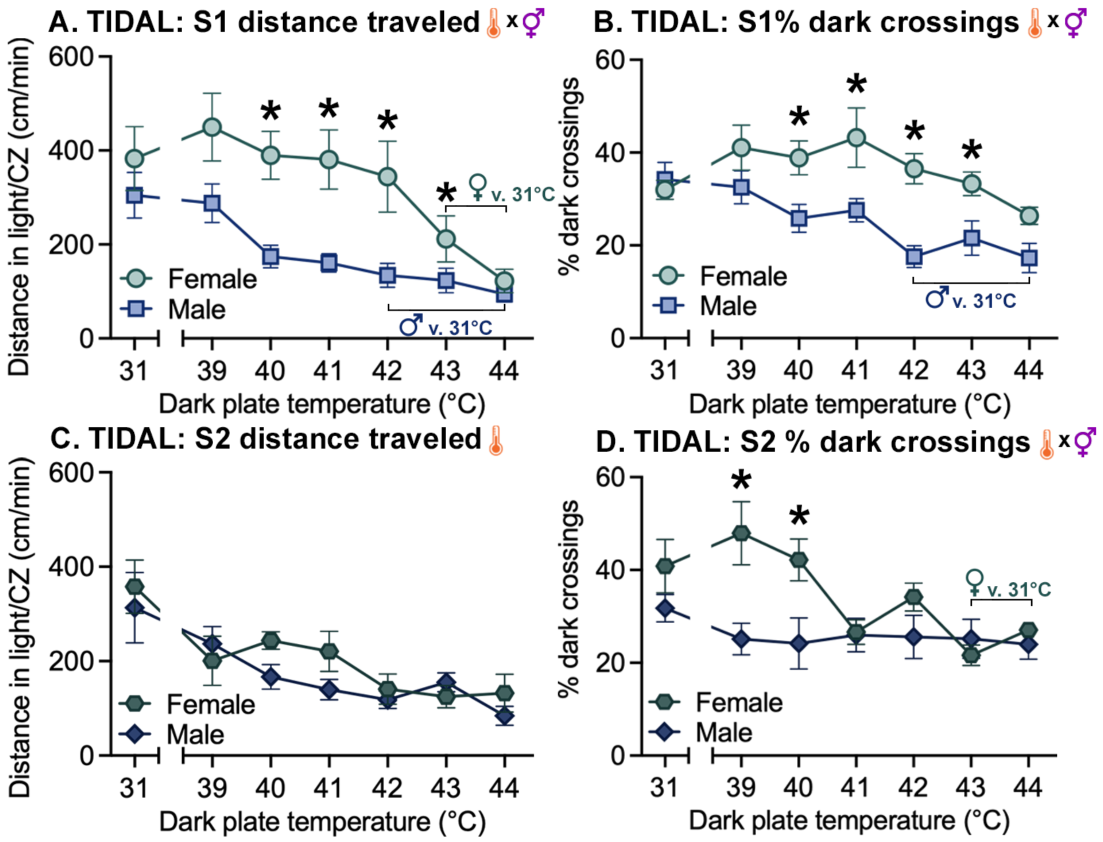


**Figure S3.** Distance traveled, percent dark crossings, and dark plate preference including time spent in center zone in our initial TIDAL experiment with optimized temperature range. **A.** In the TIDAL test, females traveled further per minute in the illuminated area compared to males as temperatures increased. **B.** Females had higher preference for crossing into the dark chamber as temperatures increased. **C.** Female (vs. male) mice showed increased dark plate preference from 39-44°C. **D.** In a second session of the TIDAL test, distance traveled was similar across sexes.

* indicates $p < 0.05$ between female and male mice; "thermometer x gender" symbol indicates significant temperature x sex interaction; "female/male symbol v. 31°C" indicates within-sex difference vs. baseline 31°C.

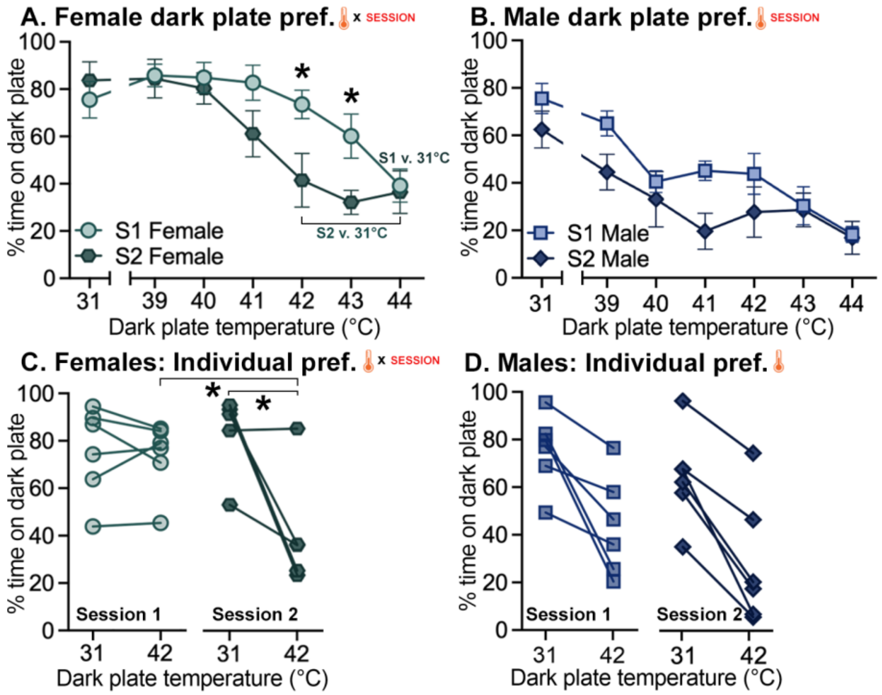


**Figure S4.** Over two sessions of TIDAL conflict testing, female (left panels) and male (right panels) mice exhibit signs of learning by avoiding the heating-dark side more quickly. **A, B.** In the TIDAL conflict test, mice of both sexes in Session 2 (vs. Session 1) showed expedited decreases in preference for the heated-dark plate. **C, D.** Session 1 and 2 dark plate preferences of individual mice with the dark plate at 31°C and 42°C. * indicates $p < 0.05$ between female and male mice; "thermometer x SESSION" symbol indicates significant temperature x session interaction; thermometer symbol alone indicates significant main effect of temperature.

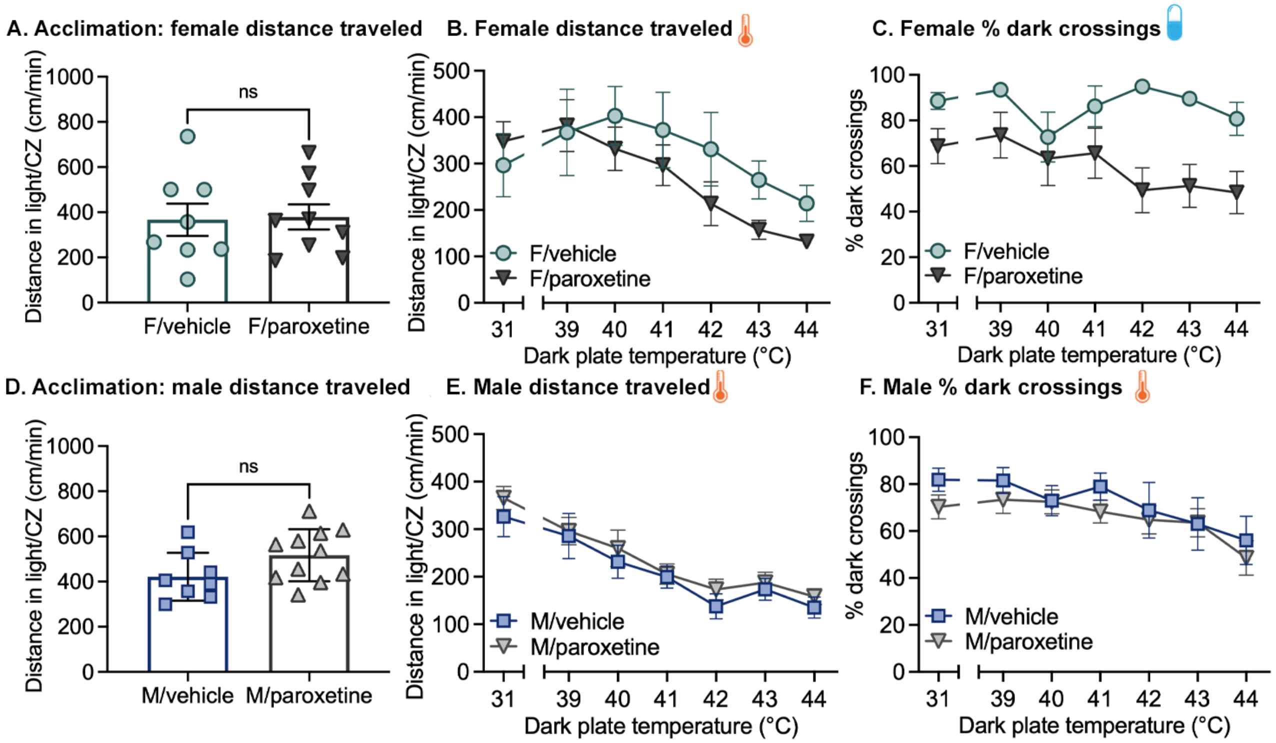


**Figure S5.** Acclimation distance traveled, overall distance traveled, percent dark crossings, and dark plate preference including time spent in center zone throughout the TIDAL conflict test after administration of paroxetine or vehicle control. **A,D.** Females and males travelled similar distances in the illuminated area in the acclimation period (dark-light test) regardless of drug. **B,E.** Mice of both sexes reduced movement with time; females traveled more per minute with the heating-dark plate. **C.** Vehicle females crossed more frequently into the dark-heated plate relative to paroxetine females. **F.** Male mice exhibited similar number of crossings into the dark plate regardless paroxetine administration.

* indicates $p < 0.05$ between F/paroxetine and F/vehicle mice; “thermometer x pill” symbol indicates significant temperature x drug interaction; thermometer or pill symbols alone indicate significant main effects of temperature and drug, respectively. Parox. v. 31°C indicates a within-group difference vs. baseline 31°C.
